# Turgor “limitation” of tree height: no evidence from leaf epidermal cell anatomy

**DOI:** 10.1101/2024.10.06.616874

**Authors:** G. Alemán-Sancheschúlz, M. E. Olson, J. A. Rosell, I. H. Salgado-Ugarte, A. I. Pérez-Maussán

## Abstract

**Highlight:** Leaf cell size distributions failed to show key signatures of the turgor limitation of tree height; instead, it is more likely that natural selection favors the cell sizes plants display.

The turgor limitation hypothesis (TLH) predicts that reduced turgor caused by gravity and hydraulic resistance at the treetop makes cell expansion beyond a certain size impossible. This environmentally-imposed upper limit on cell size should be manifest as an abrupt right-hand threshold in cell size distributions. Selection is “limited” from reaching the favored cell size, which remains on the right-hand, inaccessible, side of the developmental threshold. The TLH thus predicts that as trees grow taller, treetop leaf cell size distributions should become increasingly narrow–as selection pushes as close as possible toward the inaccessible, favored values– with a right-hand threshold. To test this, we sampled 58 individuals of two tree species across different heights, along with 24 canopy-dominant rainforest species. We measured guard, subsidiary, and pavement cell lengths from apicalmost leaves. Contrary to TLH predictions, leaf cell dimension distributions were symmetrical, without right-hand thresholds, even in the tallest trees. Regression analyses mostly revealed positive or non-significant relationships between cell size, skewness, variance, and tree height, rather than the predicted negative trends. Our findings suggest that, instead of turgor-imposed “limitation,” it is more plausible that trees produce the heights, leaf sizes, and cell dimensions favored by selection in a given environment.

## Introduction

Trees vary conspicuously in their maximum heights across habitats. Trees are tallest in relatively cool habitats that are moist and humid year-round (Givnish *et al*., 2014; Kunwar *et al*., 2021; Smith *et al*., 2023; Larjavaara *et al*., 2024). In places with higher temperatures, lower soil moisture, and higher vapor pressure deficit, tree heights are predictably lower (Tao et al., 2016; Fang *et al*., 2021; López *et al*., 2021; Flo *et al*., 2022; Chen *et al*., 2023; Smith *et al*., 2023).

These trends reach their conspicuous minima in deserts and shrublands, where trees are entirely absent, with the woody vegetation represented by shrubs and subshrubs (Liu *et al*., 2019; Tumber-Dávila *et al*., 2022). Moreover, even within a single tree species, individuals can exhibit variation in height due to environmental changes at the same location. When moist habitats experience drought, it is very common to observe terminal dieback (Rood *et al*., 2000, 2003; Walthert *et al*., 2021). The apical branches of trees die, and the tree resprouts from lower branches, resulting in a shorter tree (Davis *et al*., 2002; Koch *et al*., 2004; Camarero *et al*., 2015; Fajardo *et al*., 2019; Andrade Bueno *et al*., 2020; Kiorapostolou *et al*., 2020; Anfodillo and Olson, 2021; Olson *et al*., 2023). Understanding the causes of tree height variation and dieback is especially important given that dieback events are becoming more frequent globally as droughts become more frequent and severe due to climate change. Because so many forest properties, such as individual tree reproductive output or carbon storage, scale with tree size with slopes >>1, lowered forest stature has disproportionate impacts on forest function (Lindenmayer, 2016, 2017; Lindenmayer and Laurance, 2017; Fajardo *et al*., 2019; Enquist *et al*., 2020). It is therefore essential to understand the causes of variation in tree height and dieback.

A widely accepted hypothesis explaining tree height variation is the “turgor limitation hypothesis” (Ryan and Yoder, 1997; Koch *et al*., 2004; Woodruff *et al*., 2004; Ryan *et al*., 2006; Woodruff and Meinzer, 2011; Zhang *et al*., 2012; Fulton *et al*., 2014; Peters *et al*., 2021; Fang *et al*., 2021; Cabon and Anderegg, 2021; Potkay *et al*., 2021; Potkay and Feng, 2022; Cabon *et al*., 2024). The hypothesis builds on the foundational observation that cell expansion in plants occurs via positive turgor pressure, which overcomes the resistance of (as-yet unlignified or loosened) cell walls and the pressure exerted by surrounding tissues (Lockhart, 1965; but see Passioura and Fry 1992; Kroeger *et al*., 2011; Peters *et al*., 2021; Ali *et al*., 2023; Bou Daher *et al*., 2024). Any factor that limits turgor can therefore limit cell expansion. Gravity, for instance, affects xylem pressure with height above the ground. From the base of a tree toward the tip, xylem potential is about 0.01 MPa more negative with each meter (Woodruff *et al*., 2004). From this point of view, the upper reaches of taller trees, especially in drier areas, should have more highly negative xylem pressures.

Some correlational studies give reason to think that more negative xylem pressures can lead to lower turgor pressures (Conti *et al*., 2020; Hernández-Hernández *et al*., 2020; Epron *et al*., 2021; Knipfer *et al*., 2021; De Swaef *et al*., 2022; Oliveira *et al*., 2022; Yi *et al*., 2022; Castillo-Argaez *et al*., 2024). Soil water potentials are more highly negative in drier sites, so ground-level xylem potentials are more highly negative on drier sites. According to the turgor limitation hypothesis, drop in xylem potential due to gravity would therefore reach levels prohibitive of cell expansion at shorter heights above the ground on drier sites than on moister ones, explaining variation in tree height with moisture availability (Cabon *et al*., 2020; Hernández-Hernández *et al*., 2020; Segovia-Rivas and Olson, 2023). At the same time, if the increase in hydraulic pathlength that accompanies height growth imposes greater resistance to conductance (but see Anfodillo and Olson, 2024), then taller trees would also experience ever-lower turgor toward their upper reaches (Ryan and Yoder, 1997).

The turgor limitation hypothesis (TLH) attempts to account for a variety of phenomena, including tree height variation across habitats as well as branch shedding and lowering of stature with drought. The TLH posits that factors including soil water potential, gravity, and hydraulic pathlength resistance impose ever-more negative xylem potentials, such that cell expansion is impossible. Cessation of cell expansion necessarily stops height growth. Therefore, plants in drier areas should be shorter. If conditions in moist areas suddenly become drier, then plant height should become lower, making the TLH as outlined in the literature (Ryan and Yoder, 1997; Koch *et al*., 2004; Woodruff *et al*., 2004; Woodruff and Meinzer, 2011; Zhang *et al*., 2012; Peters *et al*., 2021; Cabon and Anderegg, 2023; Potkay *et al*., 2021) consistent with observed patterns of variation in plant height. Likewise, other authors postulate that the large-to-small gradient in leaf size from the bottoms to the tops of trees is also explained by the effects of turgor on leaf expansion (Koch *et al*., 2004; Pantin *et al*., 2011; Ding *et al*., 2020; Coussement *et al*., 2020; Long *et al*., 2020). Under these accounts, cell size, and therefore the possibility of leaf expansion, becomes smaller toward the tops of trees, tracking height-associated turgor drops.

Analogous roles are invoked for variation in height across habitats, dieback and resprouting, and conduit diameter variation along the stem and across growth rings, among other important functionally significant patterns.

As with other “limitation” hypotheses (see explorations of carbon, height, and cell size “limitation” by Körner *et al*., 1989; Körner, 2003 a, b, 2008, 2009, 2012 a, b, 2023; Körner and Hoch, 2023; Körner *et al*., 2023), one of the great advantages of the TLH is that it makes very precise and testable predictions. We outline key predictions here, which we then subject to testing. The TLH implies three main reasoning steps.

The first is that height growth ceases when the turgor that apical leaf and shoot cells can develop is insufficient for cell expansion. This means that the “limit” referred to by the TLH is an absolute threshold of developmental possibility. That is, given the context of soil potential, gravity, and conductive pathlength, developmental conditions are such that beyond a certain tree height it is impossible for treetop cells to expand.

In the second reasoning step, we can then ask what the TLH posits that plants are “limited” from. “Limit” implies that plants are prevented by turgor from reaching some unmet goal. Unless there is some “unmet goal,” it makes no sense to speak of “limitation” (cf. Körner *et al*., 1989; Körner, 2003 a, b, 2008, 2009, 2012 a, b, 2023; Körner and Hoch, 2023; Körner *et al*., 2023). But notions of “unmet goals” have an anthropomorphic tint that must be translated into naturalized biological terms. In this case, an “unreached goal” can only refer to a trait value favored by selection but not accessible because of the turgor-induced developmental threshold (Segovia-Rivas and Olson, 2023). By the lights of the TLH, selection would favor taller trees, larger cells, or longer leaves in a given environmental situation, but the developmental threshold imposed by insufficient turgor means that these would-be favored trait values are developmentally impossible. In this way, environmental conditions “limit” trees from reaching greater heights.

Finally, the TLH implies a very particular shape of trait value distributions (Fig. 1A). In the previous step, we have seen how the TLH implies that as trees grow taller, the distribution of tree heights at a given site, xylem conduit diameters in a stem, leaf lengths at the top of the crown, or the distribution of leaf cell sizes, must reflect the turgor-imposed threshold of developmental possibility. In the distribution of trait values, this absolute developmental threshold would take the form of an abrupt (vertical) right-hand of the trait distribution, corresponding to the maximum trait value permitted by turgor. What is more, the trait distribution should be extremely narrow. This is because selection should push trait values as close as possible to the threshold of developmental possibility, given that the would-be favored value lies on the other side of the threshold. The TLH thus predicts a highly distinctive trait distribution, making the hypothesis readily testable.

**Figure 1.**
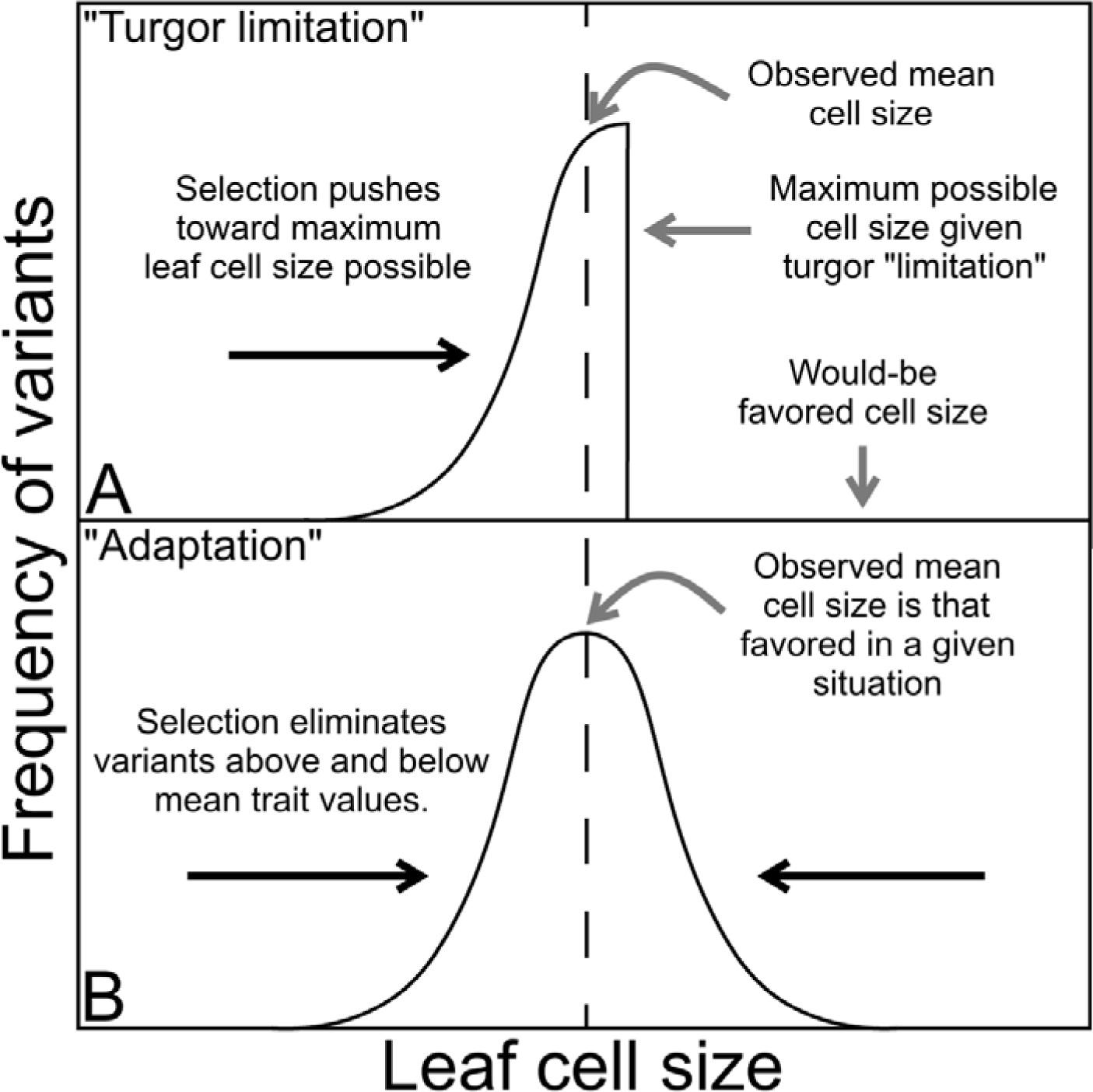
The turgor limitation hypothesis (TLH) and our “adaptation” alternative make clear and readily tested predictions. A) The TLH implies that the effects of soil potential, gravity, and path length impose an upper limit to leaf cell expansion, represented by the abrupt vertical threshold on the right-hand side of the distribution. “Limitation” implies an unreached goal; in biological terms, this means that selection is unable to reach the would-be favored trait value, which remains on the right side of the developmental threshold. Since selection pushes populations as close as possible to the threshold of ontogenetic possibility, the TLH would result in a distribution with a strong negative skewness and low variance. B) According to our adaptationist hypothesis, individuals with larger or smaller leaf cells are possible, but these individuals would have lower fitness than those individuals with cells closer to the average. In this scenario, the distribution of leaf cell length would be more normal-like or with slight positive or negative skewness, and with a greater variance than in TLH scenario, certainly without an abrupt right-hand threshold. See also Segovia-Rivas and Olson (2023).

We tested the TLH by studying the length distributions of three cell types that make up the leaf epidermis, which were guard, subsidiary, and pavement cells (also called ordinary epidermal cells, Mauseth, 1988). We used three different and complementary approaches. The first approach examined leaf cell size distributions from seedlings to trees of maximum height at a given site in *Bursera simaruba*. In the second approach we explored leaf cell dimensions from small to the tallest trees in a stand of *Eucalyptus camaldulensis* experiencing drought-induced dieback. For the third approach, we used trees of different species that make up the canopy of a primary tropical rainforest and therefore at least some individuals must be close to the maximum possible height at the study site. We provide detailed rationales regarding our three sampling strategies in the methods section. In no case did we find evidence of the distribution predicted by the TLH. We also did not find evidence of a decrease in cell size in treetop leaves from seedlings to tall plants, or decreases in cell length variances. In the light of these results, we discuss a more plausible alternative hypothesis, which is that plant cell dimensions are not “limited” in the way that the TLH depicts. Instead, plants can produce a wide array of cell lengths (or other traits, such as plant heights or leaf lengths) at any given site, but that selection favors certain values, and these are the commonly observed ones (Fig. 1B).

## Materials and methods

According to the turgor limitation hypothesis (TLH), if a tree is close to or at its height limit given its environmental conditions, leaf cells should show evidence of inability to expand given turgor drops with height (Potkay *et al*., 2021). The hypothesis implies that these environmental conditions “limit” the plant from being able to produce the cell dimensions that natural selection would favor if they were possible (Segovia-Rivas and Olson, 2023). Selection therefore should favor variants that can produce cell dimensions as close as possible to the favored values.

Consequently, the TLH predicts a very distinctive frequency distribution of cell sizes, with a strong negative skew and low variance (i.e. very narrow and asymmetrical, with the left tail of the distribution longer than the right; Fig. 1A). This pattern is so distinctive that it should be readily detectable in the data. We used three sampling strategies to test the predictions of the TLH, using frequency distributions of leaf epidermal cell dimensions.

### Seedlings to maximum tree size within a species

The first strategy involved measuring the distribution of leaf epidermal cell dimensions in different individuals of the same species, from seedlings to the tallest individuals at our study site. If the TLH is correct, then a shift in epidermal cell dimension frequency distributions should be evident from short to tall trees. The hypothesis implies that at small sizes, plants are well below the critical height at which turgor “limitation” to cell expansion would occur. Therefore, small plants can produce the mean cell dimensions favored by natural selection, given environmental conditions. Because small plants are below the height at which a turgor-imposed upper limit to cell expansion occurs, a more symmetrical distribution than that depicted in Fig. 1A should be observed. As trees grow and they approach the developmental threshold, the more symmetrical distribution should shift toward a very narrow one that is abruptly cut off at its upper limit, that is, a negatively skewed distribution with a very low variance, with a limit imposed by cell turgor.

Although this prediction is straightforward, it is challenging to find a system in which to test it. In dense forests, plants often shift from low leaf mass per unit area (LMA) shade leaves to high LMA sun leaves (Martin *et al*., 2020) as they grow from the moist, cool, and shady understory to the drier, warmer, and sunny canopy. Low LMA leaves can be achieved by larger cells, with less cell wall material per unit volume, and high LMA leaves by smaller cells, which lead to more cell wall material per unit volume (Poorter *et al*., 2009; Coble and Cavaleri, 2015). So, natural selection favoring a shift from low to high LMA with height growth could lead to a shift from large to small leaf epidermal cells. This process has nothing to do with turgor “limitation” but parallels part of the expected pattern, a decrease in cell size with height.

Therefore, we selected a species that is uniformly exposed to sun across all size classes. On the coastal plain near the Los Tuxtlas field station, Veracruz, Mexico, *Bursera simaruba* (L.) Sarg. (*Burseraceae*, *Sapindales*) is used as a living fence in cattle pastures. This means that the terrain is maximally open and sunny, with even small individuals exposed to full sun. The living fence posts are about 1.5 m tall, with branches of varying heights on individuals that have escaped pruning. Furthermore, occasional individuals in abandoned fence lines reach more than 25 meters, and seedlings often establish around the fences and along roads, providing an abundance of seedlings <1.5 m tall, also exposed to full sun. This situation provided the full range of sizes from seedlings to the largest plants at the site all with fully exposed “sun” leaves. We collected apicalmost (uppermost in the treetop) leaves from 29 individuals of *B*. *simaruba* ranging in height from 14.5 cm to 26.6 m.

### Trees experiencing dieback

The TLH would also account for tree dieback, because drought should make xylem potential more highly negative, lower the turgor that cells can develop at their apices, and therefore lower the maximum developmentally possible tree height at a given site, resulting in the sacrifice of apical branches and resprouting from lower ones (cf. Davis *et al*., 2002; Fang *et al*., 2021). Signs of turgor limitation, as in Fig. 1A, should be very clear in trees experiencing dieback because they are, according to the TLH, manifestly at their height limits. Therefore, our second strategy involved sampling a population of *Eucalyptus camaldulensis* Dehnh. (*Myrtaceae*, *Myrtales*) experiencing dieback.

University City, of the National Autonomous University of Mexico (UNAM), is built on a 2000 year old lava field, with practically no soil accumulation. The climate is characterized by a summer rainy season that receives about 1 meter of rainfall, and a six-month dry season. Given the climatic and edaphic conditions, the natural vegetation is a xeric shrubland with only very short trees in moist lava crevices (Rzedowski, 1990). *Eucalyptus camaldulensis* escaped from cultivation and is invasive on large parts of the remnant natural vegetation. Recent severe droughts (Ortega-Gaucin *et al*., 2016; CONAGUA, 2024) have resulted in marked dieback and resprouting of most of the larger individuals. As we did with *Bursera simaruba*, we sampled 29 individuals of *Eucalyptus camaldulensis*, with heights ranging from 1.09 m to 23 m. All of the large individuals showed evidence of dieback in the form of dead uppermost branches (Supplementary Fig. S1).

We collected the apicalmost leaves in each tree across all three sampling strategies. For this purpose, we used either clippers or pruner poles for trees less than 8 meters tall, or a DeLeaves canopy sampling system (Outreach Robotics, Quebec, Canada) carried by a Matrice 300 RTK drone (DJI, Shenzhen, China) (Supplementary Fig. S2). We measured tree heights using a measuring tape, and for the tallest trees, we used a Mini 4 Pro drone (DJI, Shenzhen, China) with the onboard barometric altimeter. We reset the “0” height value of the drone at the tree base, set the camera gimbal horizontal, and measured the tree height using the crosshairs on the controller screen. We repeated this process three times for each tree and averaged the measurements to determine the tree height.

### Canopy trees across species

The TLH posits that trees grow to the height permitted by the environmental conditions prevailing at any given site (Woodruff *et al*., 2004). Therefore, our final sampling strategy focused on trees of maximum height in the tropical rainforest at the Los Tuxtlas field station, Veracruz, Mexico. The station protects primary rainforest that has never been logged. This forest is subject to a marked late-winter dry season when temperatures routinely exceed 30°C. Summer-fall is characterized by heavy rainfall. Soils are relatively recent volcanic and very well-drained (Flores-Delgadillo *et al*., 1999). This combination leads to a canopy that is uniformly about 30 m tall. No increase or decrease in the mean forest height has been observed since the field station was established. The TLH depicts the effects of environmental “limitation” being pervasive, even affecting cell dimensions along the lengths of stems in all habitats (Cabon and Anderegg, 2021). Therefore, if the TLH is correct, it is reasonable to expect that at least some of the canopy individuals are at or near their maximum heights permitted by environmental conditions via their effects on leaf cell turgor and that they should show at least some evidence of the very distinctive cell size distribution.

We sampled apicalmost leaves from the following 24 species from 12 families: *Alchornea latifolia* Sw. (*Euphorbiaceae*, *Malpighiales*), *Bernoullia flammea* Oliv. (*Malvaceae*, *Malvales*), *Ceiba pentandra* (L.) Gaertn. (*Malvaceae*, *Malvales*), *Clarisia biflora* Ruiz & Pav. (*Moraceae*, *Rosales*), *Cojoba arborea* (L.) Britton & Rose (*Fabaceae*, *Fabales*), *Cordia megalantha* S. F. Blake (*Cordiaceae*, *Boraginales*), *Cordia stellifera* I. M. Johnst. (*Cordiaceae*, *Boraginales*), *Cupania glabra* Sw. (*Sapindaceae*, *Sapindales*), *Damburneya ambigens* (S.F. Blake) Trofimov (*Lauraceae*, *Laurales*), *Dendropanax arboreus* (L.) Decne. & Planch. (*Araliaceae*, *Apiales*), *Ficus aurea* Nutt. (*Moraceae*, *Rosales*), *Ficus yoponensis* Desv. (*Moraceae*, *Rosales*), *Inga acrocephala* Steud. (*Fabaceae*, *Fabales*), *Lonchocarpus guatemalensis* Benth. (*Fabaceae*, *Fabales*), *Ocotea uxpanapana* T. Wendt & van der Werff (*Lauraceae*, *Laurales*), *Platymiscium dimorphandrum* Donn. Sm. (*Fabaceae*, *Fabales*), *Poulsenia armata* (Miq.) Standl. (*Moraceae*, *Rosales*), *Pterocarpus rohrii* Vahl (*Fabaceae*, *Fabales*), *Sapium lateriflorum* Hemsl. (*Euphorbiaceae*, *Malpighiales*), *Sideroxylon portoricense* Urb. (*Sapotaceae*, *Ericales*), *Ulmus mexicana* (Liebm.) Planch. (*Ulmaceae*, *Rosales*), *Vatairea lundellii* (Standl.) Killip ex Record (*Fabaceae*, *Fabales*), *Vochysia guatemalensis* Donn. Sm. (*Vochysiaceae*, *Myrtales*), and *Zanthoxylum riedelianum* Engl. (*Rutaceae*, *Sapindales*).

### Anatomical methods and measurement of quantitative variables

We pickled leaves in 70% ethanol. Subsequently, we made stomatal impressions using the nail polish method (Wu and Zhao, 2017). We then mounted these impressions on slides, observed them under an NS200 compound microscope (Nachet, Paris, France), and photographed them with an A6000 digital camera (A6000, Sony Corp., Tokyo, Japan). From the stomatal impressions, we measured the lengths of three leaf cell types, guard, subsidiary, and pavement, 100 cells each, using ImageJ 1.53k (National Institutes of Health, Maryland, U.S.A.), calibrating the images with a Nikon stage micrometer (MBM13100, Nikon Corp., Tokyo, Japan), to convert cell length measurements from pixels to micrometers.

### Statistical analyses

First we log_10_ transformed the lengths of guard, subsidiary, and pavement cells (Kerkhoff and Enquist, 2009). Cell dimensions not only result from multiplicative growth processes, but the metabolic and biomechanical consequences of the same absolute size increment are very different at different points along the cell size range. For instance, a 5 µm difference between cells that are 25 and 30 µm in diameter has much more significant consequences than the same 5 µm difference between cells that are 255 and 300 µm in diameter. In smaller cells, a 5 µm difference represents a substantial proportion of the total cell diameter, leading to significant changes in surface area, volume, and, consequently, the mechanical and metabolic properties of the cell. Mechanically, smaller cells experience a greater relative change in their structural integrity, wall tension, and susceptibility to external forces when their diameter changes by 5 µm. Metabolically, a small cell’s resource allocation, nutrient uptake, and metabolic rates can also be heavily influenced by such a change, given that the cell’s volume increases more sharply in proportion to its surface area. In contrast, for much larger cells, a 5 µm difference represents a nearly negligible fraction of the total diameter, resulting in relatively minor changes to the cell’s overall biomechanical and metabolic characteristics. The mechanical impact, such as changes in wall stress or structural stability, is less pronounced because the larger cells already have more substantial structural support. Similarly, the metabolic consequences are less significant, as the change in volume relative to surface area is less dramatic, leading to more modest adjustments in nutrient uptake and metabolic activity. Log transformation is appropriate in these situations because it effectively scales the data in a way that makes these proportional and functionally significant differences comparable.

We then inspected the distributions of leaf cell length across all individuals using kernel density estimators and weighted averages of rounded points. Kernel density estimators are a non-parametric approach that depict cell length distributions as smoothed versions histograms (Salgado-Ugarte *et al*., 1993). This method offers significant advantages over traditional histograms, such as independence from interval selection and greater adaptability to the underlying data structure. Additionally, it can reveal important characteristics of the data, including skewness, tail weight, and the presence of multiple modes (Silverman, 1986). The parameters of this method included the Gaussian kernel, Silverman’s optimal bandwidth and the average of 40 histograms (García-Cervigón *et al*., 2020; Salgado-Ugarte and Saito-Quezada, 2020, 2022). The weighted averages of rounded points method is used to reduce variability in estimates of discrete data or to smooth categorical data. This technique approximates a specific type of kernel density estimator (in this case, the Gaussian kernel) by assigning weights to the data points (Salgado-Ugarte *et al*., 1995). The weights correspond to the number of histograms being averaged—in this case, 40 histograms (Härdle, 1991). Using smoothed distributions is advantageous for our study because they should make detection of any right-hand threshold more immediate than using histograms with discrete bins. If the TLH is correct, then we should find evidence of the distribution in Fig. 1A. In *Bursera simaruba* and *Eucalyptus camaldulensis,* there should be a dramatic shift in the distributions of guard, subsidiary, and pavement cells length, from symmetrical (or at least without a conspicuous right-hand abrupt threshold) in small plants, to very narrow distributions with abrupt right-hand thresholds in taller plants as they approach the maximum permitted height at a given site. Kernel density estimators and weighted averages of rounded points procedures were performed in Stata v.14.1 (StataCorp, 2016).

A further way of examining the predictions of the TLH involved testing for changes in the skewness and variance of the frequency distributions of guard, subsidiary, and pavement cell lengths. As plants grow in height, the cells of the leaves are expected to become smaller due to the “limitation” imposed by cell turgor. Statistically this means that the frequency distributions should become significantly more negatively skewed. A negative skew means that most of the cell size measurements would be concentrated toward the right side of the distribution, as selection “bunches up” cell sizes against the right-hand threshold. Therefore, the TLH predicts that the skewness should shift from values near zero to negative values as the plants grow taller. At the same time, the TLH predicts a shift in the shape of the leaf cell size distribution, from broad to very narrow because of selection favoring results as close as possible to the would-be favored cell length to the right of the developmental threshold. This means that there should be a reduction in variance, which should decrease as the plants increase in height.

According to the aforementioned points, a distribution that simultaneously exhibits a strong negative skewness and low variance would have most of its values clustered toward the far right, a tail extending to the left, and a peak that is higher and narrower compared to a normal distribution (as in Fig. 1A). This necessarily implies that there should be a negative relationship between skewness and tree height, as well as between variance and tree height. Therefore, we conducted simple linear regression analyses to explore the relationship between the skewness and variances of leaf cell lengths, and tree height of *B*. *simaruba* and *E*. *camaldulensis*. These regressions were performed in R 4.4.1 (R Development Core Team, 2024).

The turgor limitation hypothesis also predicts that there should be a significant negative relationship between cell size and tree height. This is because according to the TLH, water potential becomes more negative with increase in tree height (Woodruff *et al*., 2004), and simultaneously resistance to conductance increases, so the TLH envisions taller trees experiencing lower cell turgor (Ryan and Yoder, 1997) and reduced cell expansion (Lockhart, 1965) at their upper reaches as a consequence. As plants grow taller and the turgor that can be developed drops, cell size should become markedly smaller until it reaches the minimum possible for height growth. The TLH thus predicts a significant negative relationship between epidermal cell size and height. To test this prediction, we analyzed the relationship between the mean lengths of guard, subsidiary, and pavement cells and tree height using simple linear regressions implemented through the *lm* command in R 4.4.1.

## Results

The turgor limitation hypothesis makes clear predictions regarding the very distinctive pattern of trait frequency distributions that should be observed in leaves at the apices of trees as they approach their height limits, as in Fig. 1A.

Our analyses did not find evidence consistent with the predictions of the TLH. First, inspection of the distributions of lengths of leaf guard, subsidiary, and pavement cells revealed no evidence of the very distinctive predicted frequency distribution, with its conspicuous right-hand threshold and its narrowness (Figs. 2-4, and Supplementary Figs. S3-S11). In all cases, the frequency distributions were more or less symmetrical, with most values concentrated in the center of the distribution. As is typical in the frequency distributions of biological variables, there was some variation in symmetry, variance, and apparent number of modes across leaves from individuals of different heights and species. However, none showed any evidence of a right-hand threshold, not even in the tallest *Bursera simaruba* plants (Figs. 2, and Supplementary Figs. S3-S5) or in *Eucalyptus camaldulensis* subject to dieback, which did not display the expected shift consistent with turgor limitation (Figs. 3, and Supplementary Figs. S6-S8).

**Figure 2.**
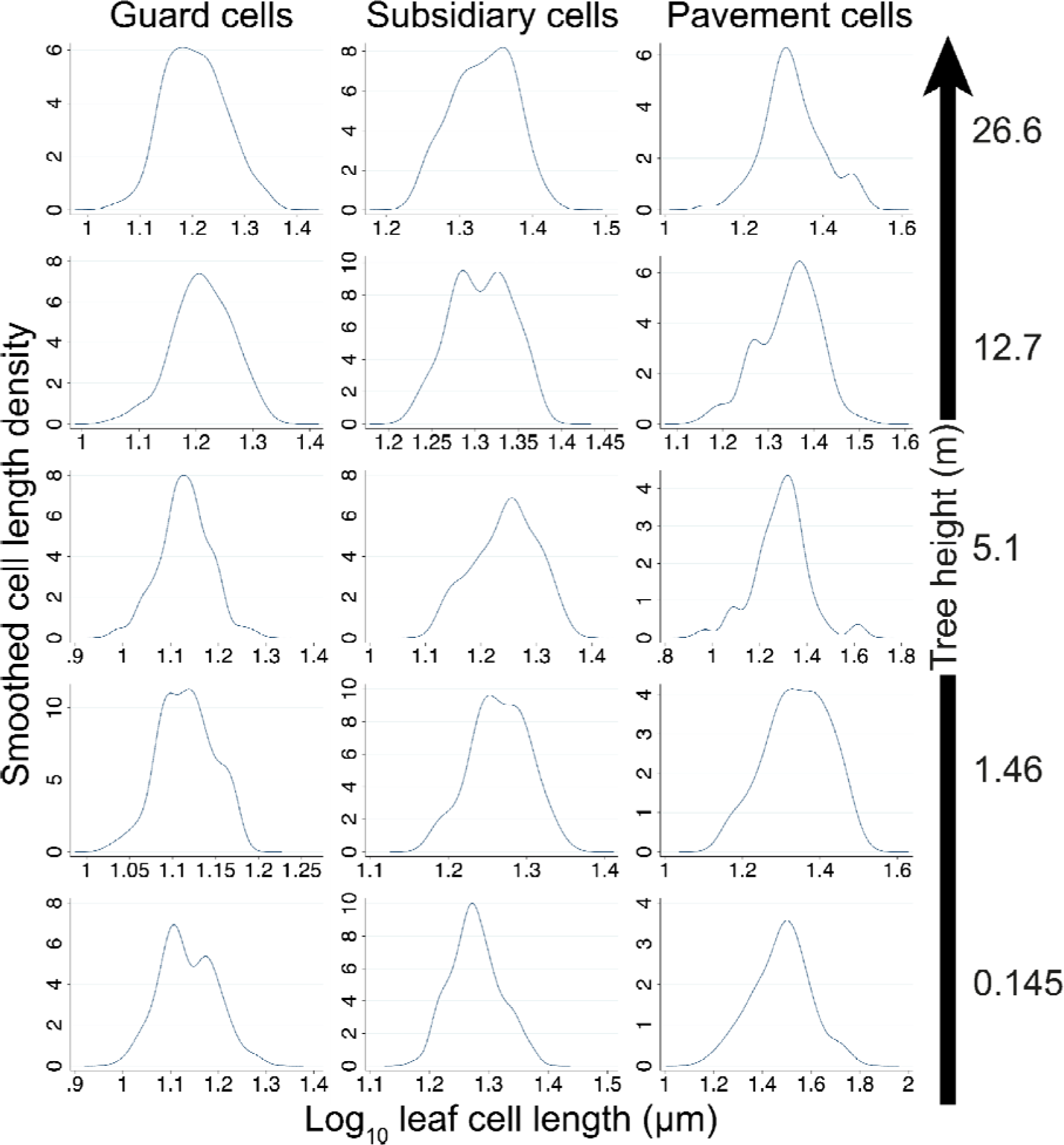
No evidence of the predicted shift in leaf cell size distribution with height due to turgor “limitation” from short to tall sun-exposed *Bursera simaruba*. The plots give representative distributions of guard, subsidiary, and pavement cell length in five representative *Bursera simaruba* individuals, from short plants at the bottom and tall plants at the top (tree height given at right). The TLH implies that frequency distributions should be more symmetrical in shorter plants, at the bottom of the figure, and that taller plants should show evidence of the right-hand threshold of developmental possibility imposed by turgor limitation. Instead, the plots depict normal-like distributions that are similar across individuals of different heights, without right-hand turgor-imposed thresholds, contradicting the predictions of the TLH. Tree height shown at right.

**Figure 3.**
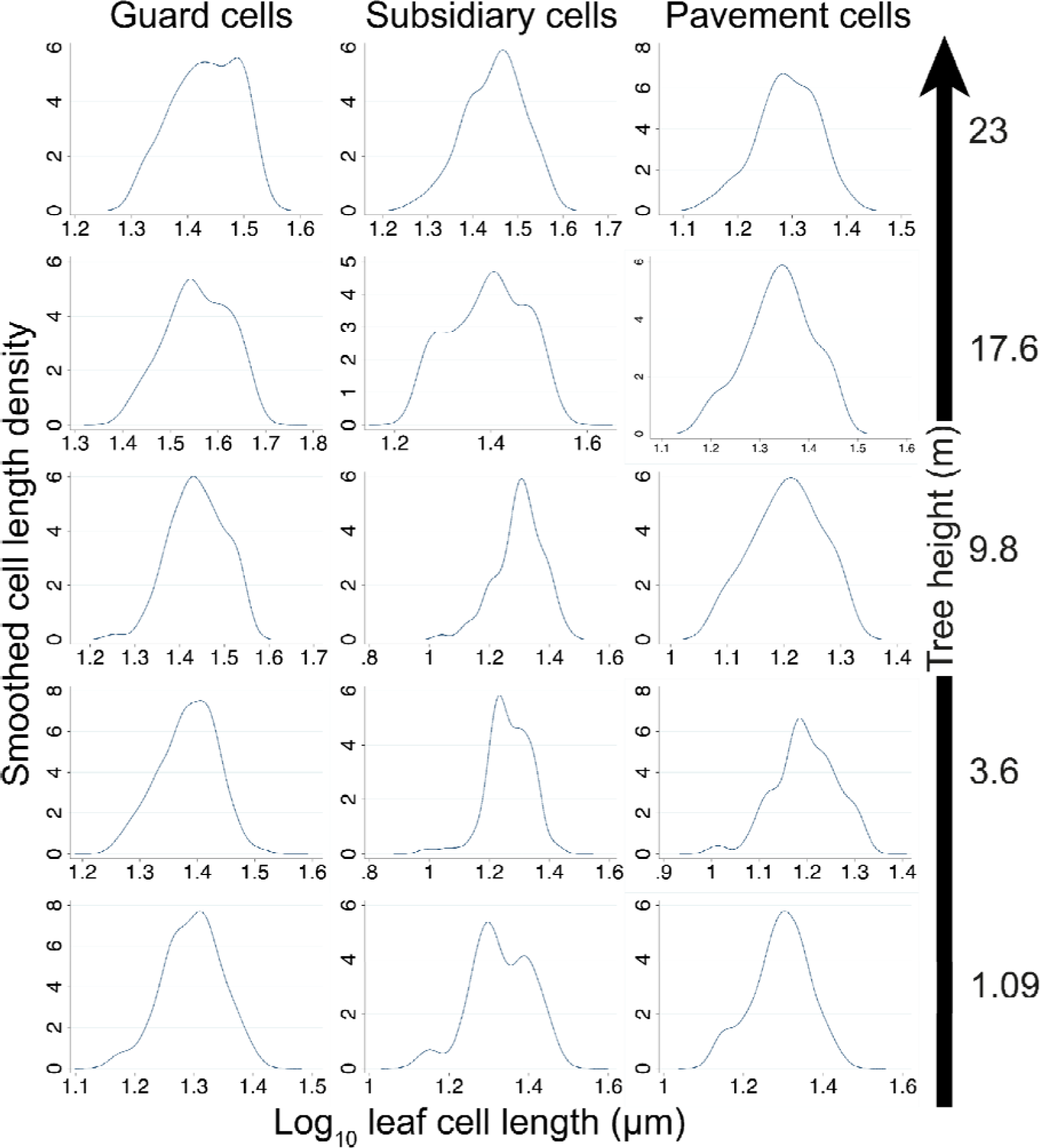
No evidence of the predicted change in leaf cell length distribution due to turgor limitation as trees grow taller and experience dieback in *Eucalyptus camaldulensis*. The plots represent the distributions of guard, subsidiary, and pavement cell length in five representative *Eucalyptus camaldulensis* individuals. The data show relatively symmetrical distributions for the five individuals. Some distributions are narrower, but the expected clustering of measurements toward the right side of the plots, predicted by the TLH, is absent. Even the tallest individual (23 m) experiencing dieback shows distributions similar to those of the shortest tree (1.09 m). Tree height shown at right.

**Figure 4.**
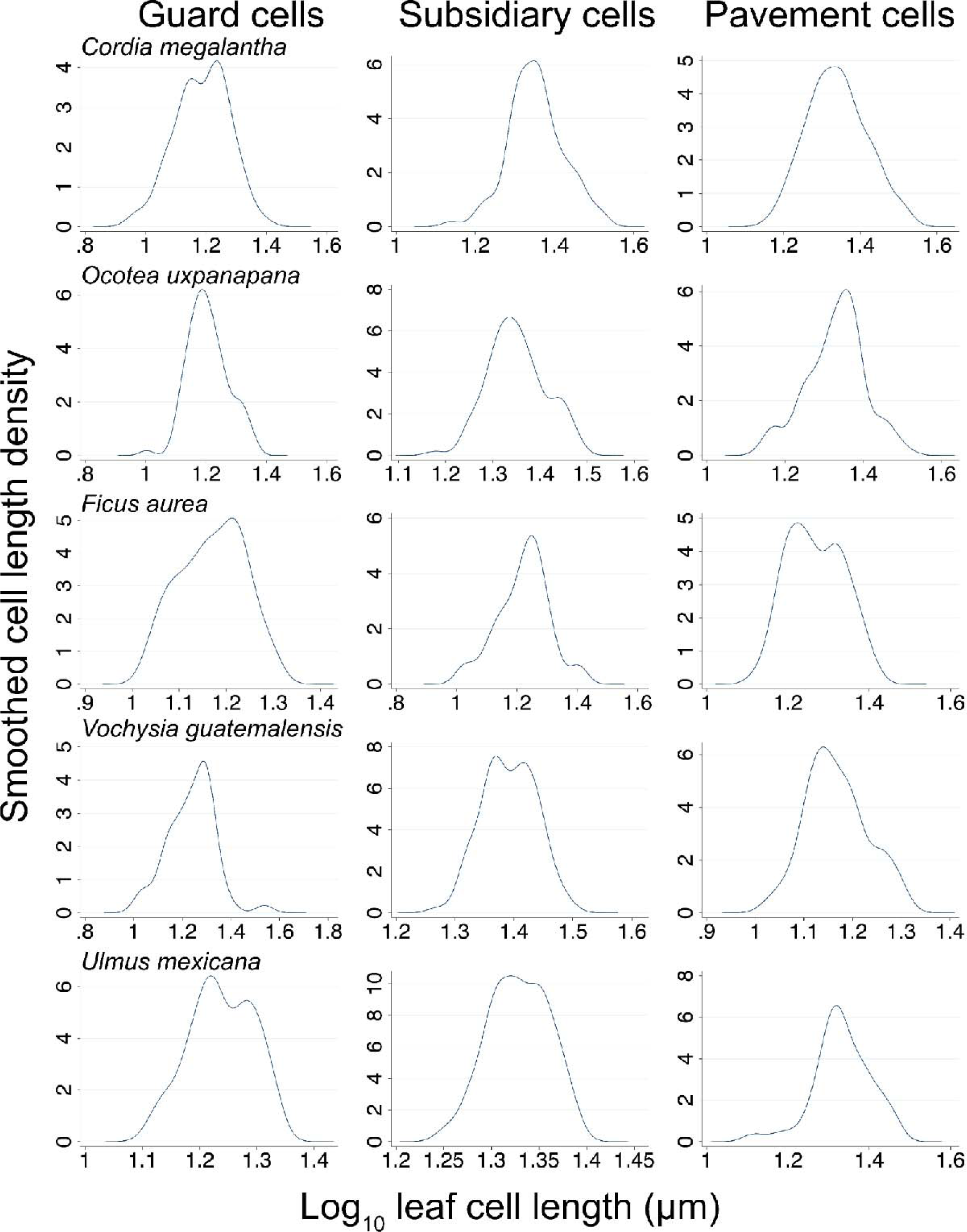
No evidence of the right-hand developmental threshold and narrow distribution of leaf predicted by the turgor limitation hypothesis in dominant forest canopy trees. The plots represent the distribution of guard, subsidiary, and epidermal cells in five representative species of the 24 measured (see Figs. S6-S8) that make up the canopy of a tropical rainforest and therefore are at or close to the maximum height for these species at our sampling site.

Likewise, if turgor “limitation” is as pervasive as it is postulated to be (Cabon *et al*., 2020) one would expect that at least one of the dominant rainforest canopy trees would show some signs consistent with it, but not a single individual showed any such evidence (Figs. 4, and Supplementary Figs. S9-S11).

Similarly, our results in *Bursera simaruba* and *Eucalyptus camaldulensis* contradicted the expectation that the distributions of leaf cell lengths should become negatively skewed with height growth. Regression analyses did not find the predicted relationships between the skewness of guard, subsidiary, and pavement cell length and the heights of trees in *B*. *simaruba* (guard cells: F_(1, 27)_= 0.73, *P*= 0.39; subsidiary cells: F_(1, 27)_= 0.15, *P*= 0.69; pavement cells: F_(1, 27)_= 0.20, *P*= 0.65) (Fig. 5A-C). In *E*. *camaldulensis*, no significant relationships were observed between skewness of leaf cell lengths and tree height (guard cells: F_(1, 27)_= 0.46, *P*= 0.50; subsidiary cells: F_(1, 27)_= 0.94, *P*= 0.33; pavement cells: F_(1, 27)_= 0.87; *P*= 0.35) (Fig. 5D-F).

**Figure 5.**
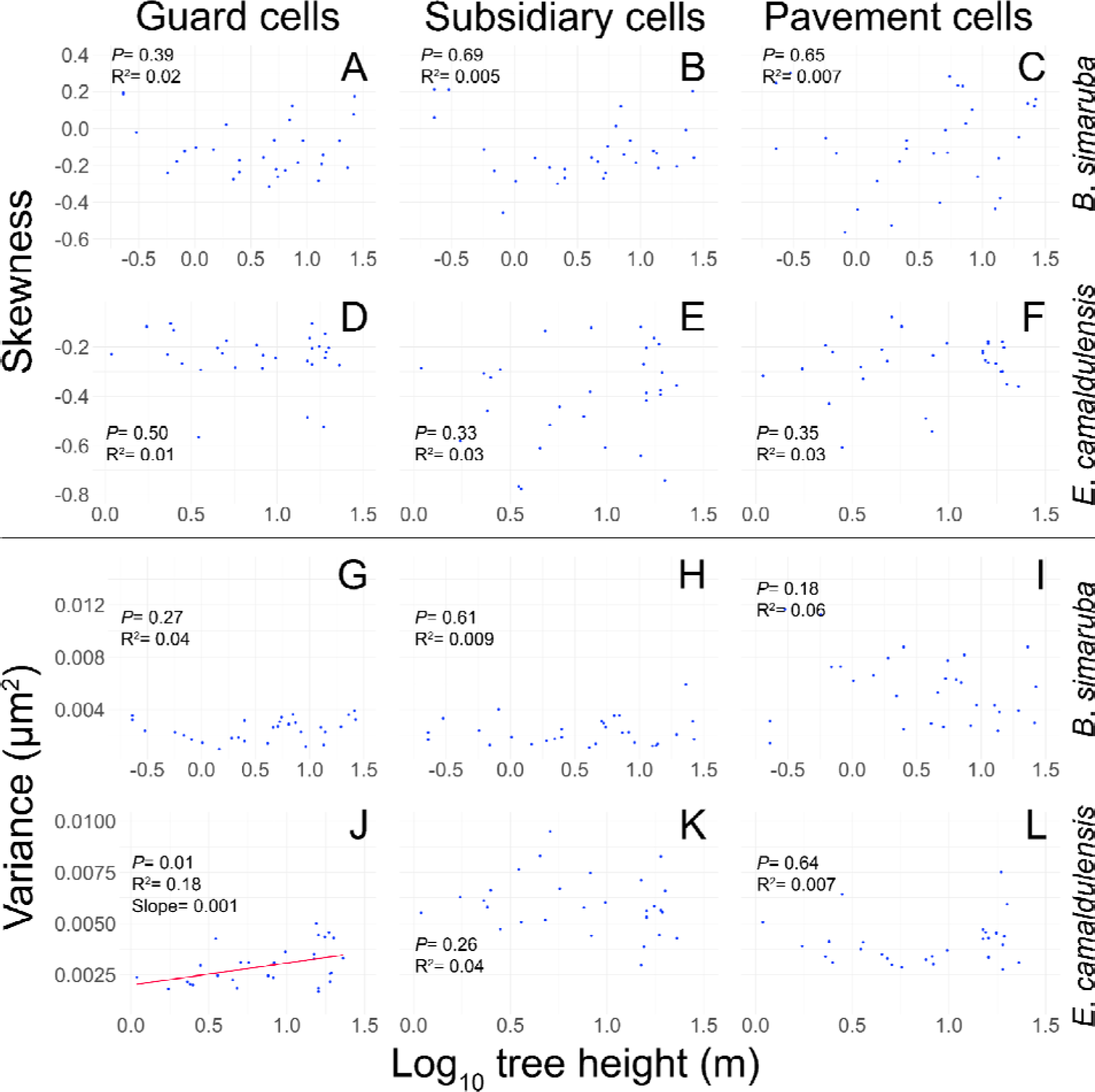
No evidence of the negative relationships between skewness and variance of leaf cell lengths, and tree height predicted by TLH. Linear regression analyses of log_10_ guard, subsidiary, and epidermal cell length skewness and variance, against log_10_ tree height, in *Bursera simaruba* (A-C, G-I) and *Eucalyptus camaldulensis* (D-F, J-L). The TLH predicts that leaf cell size distribution should exhibit a strong negative skewness and a very low variance as trees grow taller. Additionally, the skewness and variance should have a negative relationship with tree height. These data do not bear out the expected relationships between skewness, variance, and height. Scatter plots of skewness vs. height showed that the points are distributed near zero on the y-axis, indicative of distributions close to being symmetric. The variance was relatively uniform regardless of the height of the trees. The one exception were *E. camaldulensis* guard cells, which *increased* in variance with height rather than decreasing as predicted by the TLH. Each blue dot represents the skewness or variance of 100 measurements per individual, of the 29 individuals sampled per species.

Likewise, our results in *Bursera simaruba* and *Eucalyptus camaldulensis* found no evidence that the variances of leaf cell lengths should become increasingly narrow with height growth. In *B*. *simaruba*, we did not find significant relationships between the variance of leaf cells and tree height (guard cells: F_(1, 27)_= 1.23, *P*= 0.27; subsidiary cells: F_(1, 27)_= 0.25, *P*= 0.61; pavement cells: F_(1, 27)_= 1.88, *P*= 0.18) (Fig. 5G-I). Similarly, the variance of subsidiary and pavement cells did not show a relationship with tree height in *E*. *camaldulensis* (subsidiary cells: F_(1, 27)_= 1.26, *P*= 0.26; pavement cells: F_(1, 27)_= 0.21, *P*= 0.64) (Fig. 5K, L). However, we observed a significant relationship between the variance of guard cells and tree height in this species (F_(1, 27)_= 6.26, *P*= 0.01) (Fig. 5J). This relationship was positive and indicated an *increase* in variance with the growth in tree height, which is contrary to the prediction of the TLH.

Our simple linear regression analyses of mean leaf cell lengths against tree height revealed significant relationships between the lengths of the three types of leaf cells and tree height in *Bursera simaruba* and *Eucalyptus camaldulensis*. In *B*. *simaruba*, the mean lengths of guard and subsidiary cells showed positive relationships with tree height (guard cells: F_(1, 27)_= 32.95, *P*< 0.001; subsidiary cells: F_(1, 27)_= 4.76, *P*= 0.03), although subsidiary cells had a lower proportion of explained variability (Fig. 6A, B). In pavement cells, this relationship was negative, but the variances at maximum height were very similar to those across all height categories in the other two cell types (F_(1, 27)_= 14.02, *P*< 0.001) (Fig. 6C). In *E*. *camaldulensis*, significant positive relationships were observed between the mean length of the three types of leaf cells and tree height (guard cells: F_(1, 27)_= 19.88, *P*< 0.001; subsidiary cells: F_(1, 27)_= 23.59, *P*< 0.001; pavement cells: F_(1, 27)_= 7.43, *P*= 0.01) (Fig. 6D-F). In *E*. c*amaldulensis*, pavement cells showed the lowest percentage of explained variability (Fig. 6F).

**Figure 6.**
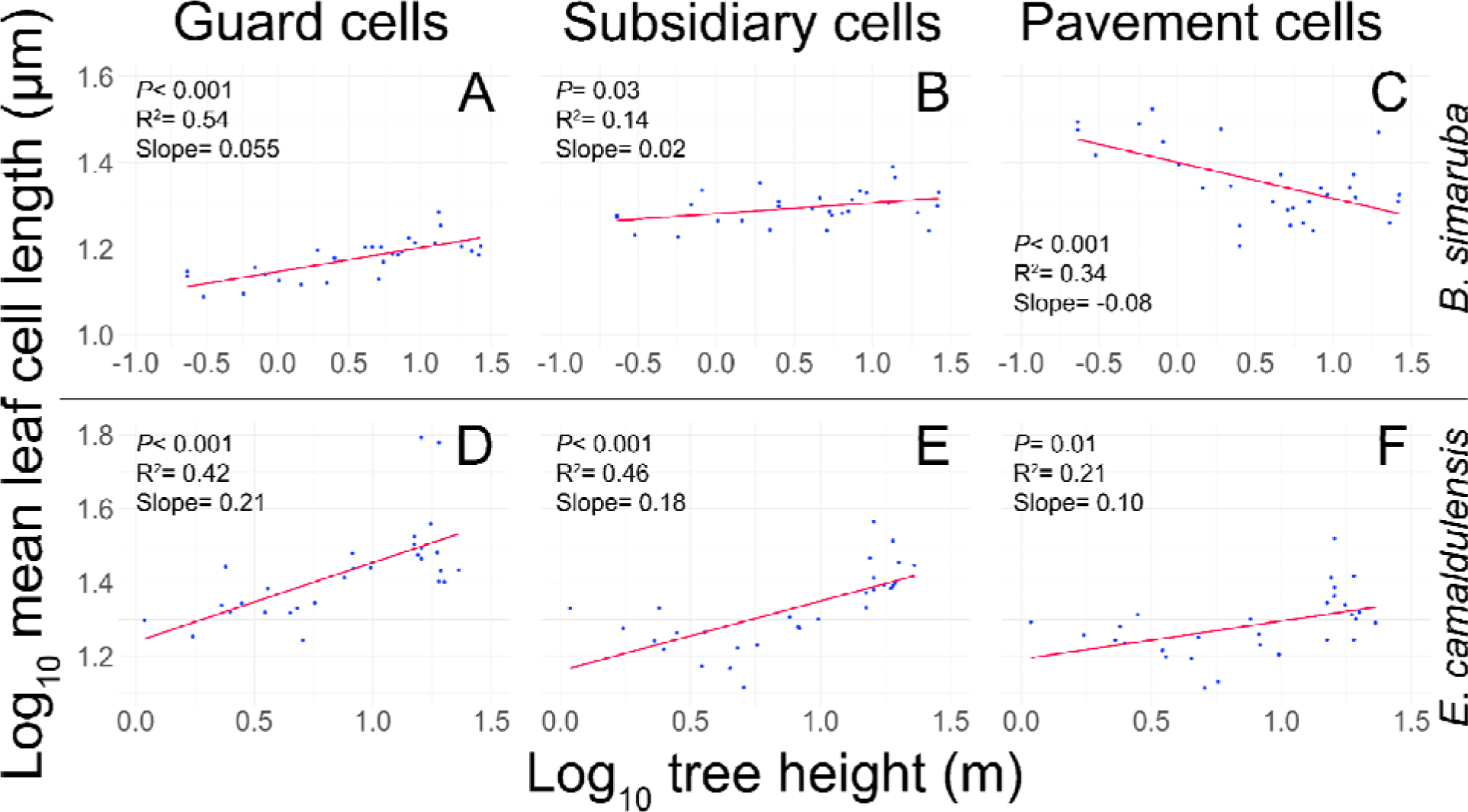
Cell size does not follow TLH predictions. The graphs show linear regression analyses of log_10_ mean leaf cell length (μm) against log_10_ tree height (m). The TLH predicts a negative relationship between leaf cell lengths and tree height, meaning that taller trees would have smaller cells. However, the regression graphs contradict the TLH, as the observed relationships are mostly positive. A-C, *Bursera simaruba*. D-F, *Eucalyptus camaldulensis*. Even in the one case where a negative relationship was observed (C, pavement cells in *B. simaruba*), this was achieved via larger cells in seedlings, with the minimum pavement cell size in large individuals being the same as the maximum value in the guard and subsidiary cells, with no butting up against some developmentally possible maximum in evidence. Each blue point represents the average of 100 measurements per individual, of the 29 sampled for each species.

## Discussion

In contrast to the predictions of the TLH, we did not find the expected shift in leaf cell length distributions from seedlings to trees with maximum height, from large to small cells with growth in tree height, or from relatively symmetrical to negatively skewed and with narrow variance (Fig. 1A). In all cases, we observed distributions that were nearly symmetrical or only slightly leaning to the left or right (skewness close to zero or somewhat positive or negative), all within the normal range of variation for biological variables. If TLH predictions were correct, there would be a negative relationship between skewness and tree height. However, in *Bursera simaruba* (Fig. 5A-C) and *Eucalyptus camaldulensis* (Fig. 5D-F), skewness of leaf cell length was unrelated to height. In fact, we are not aware of any distribution such as that predicted by the TLH ever having been observed in any plant variable (Ewers *et al*., 1990; Jacobsen *et al*., 2012, 2019; García-Cervigón *et al*., 2020; de Moraes *et al*., 2022; Silvestro *et al*., 2022; Hacke *et al*., 2017, 2023; González *et al*., 2024; Li *et al*., 2024; Matos *et al*., 2024).

Similarly, the TLH predicts that, as trees grow taller, the variance of leaf cell length should become increasingly smaller, and therefore, there should be a negative relationship between variance and tree height. Our results again contradict the predictions of the TLH, since most of the variance values were not significantly related to tree height and indicated that leaf cell size was relatively uniform among trees of different heights (Figs. 5G-I, J-L). Only the guard cells of *Eucalyptus camaldulensis* were significantly related to tree height (Fig. 5J), but this relationship was positive, meaning that the variance of guard cell length increases as trees increase in height. These findings clearly contradict the TLH predictions regarding leaf cell size variance.

Another foundational prediction of the TLH is that taller trees should have smaller leaf epidermal cells at the treetop compared to smaller individuals. However, our findings contradict this prediction. In both *Bursera simaruba* and *Eucalyptus camaldulensis*, we mostly observed a positive relationship between cell size and tree height, with larger cells in taller trees (Fig. 6).

Only in *B. simaruba* did pavement cell size negatively correlate with height (Fig. 6C). Yet, even this result contradicts the TLH, as the smallest epidermal cells at the tops of the tallest trees are still the same size as the largest guard and subsidiary cells within the same leaves and trees (Figs. 6A, B). According to the TLH, cell size should decrease with height due to a turgor-imposed developmental threshold, affecting all cell types equally. Our data, however, show no evidence of such a developmental ‘limitation’ of cell dimensions with height growth.

### What do our results mean for the turgor limitation hypothesis?

One possible response to our data is to limit the TLH’s scope, arguing that our results do not contradict it because we didn’t study trees at the global height maximum (e.g., Koch *et al*., 2004). This argument would claim that the TLH explains height limits only in trees exceeding 100 meters, not the shorter trees we examined. However, this contradicts the TLH’s broader application in the literature, which suggests it accounts for “major ecological patterns, i.e. tree growth scaling to tree size and tree maximum height in relation with water availability” (Cabon and Anderegg, 2021 p. 225, also Potkay *et al*., 2021 pp. 245-246), as well as variation in conduit diameters tip-to-base along stems and across growth rings (Cabon *et al*., 2020). Given the broad influence granted the TLH in the literature, signs of turgor limitation should have been evident in all our samples, where we studied the tallest viable trees for each species and habitat. For example, the tallest *E. camaldulensis* showed dieback and resprouting at lower heights, indicating proximity to their height limits. Our results, therefore, suggest that the TLH is either incorrect or so narrowly applicable as to be relevant only for extreme height limits. We suspect that even in the tallest trees, findings similar to ours would emerge, supporting an alternative, more plausible hypothesis.

Instead of “limitation” imposed by turgor, it is much more likely that the trait values observed at a given point on a plant are the result of natural selection rather than developmental impossibility. From this point of view, it seems almost certain that a given individual could produce larger or smaller leaf cells than the actual observed size observed at that microsite (Fig. 1B; also see Segovia-Rivas and Olson, 2023). As a result, rather than mean cell size values being the result of “limitation,” it seems more plausible that they are the product of selection in the context of a wider range of developmental possibilities. This means that for a given species, larger or smaller cells are developmentally possible, but are not favored by selection in that species. The sizes of guard cells are known to affect stomatal conductance profoundly (Lawson and Matthews, 2020), and the proportions of subsidiary cell-guard cell sizes also can profoundly affect stomatal kinetics (Rui *et al*., 2024). Likewise, epidermal cell size and shape affect leaf mechanical properties (Bidhendi *et al*., 2023). Moreover, the developmental mechanics of epidermal cell development are far more complex than simple wall relaxation-turgor mechanisms (Bidhendi *et al*., 2019). Therefore, it is virtually certain that selection finely tunes variables such as cell size rather than their being haplessly “limited” by the environment.

This adaptive interpretation aligns not only with the observations explained by the turgor limitation hypothesis (TLH) but also with our empirical data. For instance, Cabon *et al*. (2020) suggest that the tip-to-base conduit widening pattern is driven by turgor variations caused by changes in xylem water potential. While the TLH explains this pattern as a gradient in xylem water potential, it does not address why the pattern matches so closely with that predicted by evolutionary optimality models (Olson *et al*., 2020; Koçillari *et al*., 2021), which account for factors like construction cost, hydraulic resistance, and embolism risk. The striking similarity between predicted and observed patterns suggests that natural selection, rather than mere chance, is responsible. Furthermore, the TLH does not predict the patterns in our data on leaf cell length distributions (Supplementary Figs. S3-S11). From an adaptive perspective, turgor variation is part of the proximate mechanism shaping cell size, not a limitation to development imposed by external conditions (Olson, 2012; Segovia-Rivas and Olson, 2023).

## Supplementary data

The following supplementary data are available at JXB online.

**Figure S1.** *Eucalyptus camaldulensis* individual showing apical branch dieback and resrpouting at a lower height in a xeric shrubland, located in University City, of the National Autonomous University of Mexico (UNAM).

**Figure S2.** Sampling of the terminal leaves of a 21 m tall individual of *Eucalyptus camaldulensis* experiencing drought-induced dieback.

**Figure S3.** Distributions of guard cell lengths in the 29 individuals sampled of *Bursera simaruba*.

**Figure S4.** Distributions of subsidiary cell lengths in the 29 individuals sampled of *Bursera simaruba*.

**Figure S5.** Distributions of pavement cell lengths in the 29 individuals sampled of *Bursera simaruba*.

**Figure S6.** Distributions of guard cell lengths in the 29 individuals sampled of *Eucalyptus camaldulensis*.

**Figure S7.** Distributions of subsidiary cell lengths in the 29 individuals sampled of *Eucalyptus camaldulensis*.

**Figure S8.** Distributions of pavement cell lengths in the 29 individuals sampled of *Eucalyptus camaldulensis*.

**Figure S9.** Distributions of guard cell lengths in the 24 canopy dominant tree species sampled in a tropical rainforest.

**Figure S10.** Distributions of subsidiary cell lengths in the 24 canopy dominant tree species sampled in a tropical rainforest.

**Figure S11.** Distributions of pavement cell lengths in the 24 canopy dominant tree species sampled in a tropical rainforest.

## Acknowledgements

We thank Rosamond Coates and Álvaro Campos Villanueva from the Los Tuxtlas Tropical Biology Station, UNAM, for their valuable assistance in the field and laboratory work.

## Author contributions

MEO designed and conceptualized the study, and conducted fieldwork. GA-S carried out field and laboratory, and data analysis. JR and IHS-U contributed to data analysis. AIP-M conducted fieldwork. All authors participated in writing and reviewing the manuscript.

## Data availability

The data on tree heights and the lengths of guard, subsidiary, and pavement cells for all species and individuals analyzed are available in the DRYAD Digital repository, https://doi.org/10.5061/dryad.bzkh189kg (Alemán-Sancheschúlz *et al*., 2024).

